# A predatory gastrula leads to symbiosis-independent settlement in *Aiptasia*

**DOI:** 10.1101/2023.05.26.542442

**Authors:** Ira Maegele, Sebastian Rupp, Suat Özbek, Annika Guse, Elizabeth A. Hambleton, Thomas W. Holstein

## Abstract

The planulae larvae of cnidarians (jellyfish, hydras, anemones, corals) have attracted interest since Haeckels 150-year-old postulation of the gastrula developmental stage of sponges and corals as the terminal lifeform of primitive multicellular metazoans. Widely viewed as primarily particle feeders, the planulae larvae of the anemone Exaiptasia pallida (commonly Aiptasia) have not been reported to undergo settlement and metamorphosis to adult morphology, and the lack of a closed lifecycle has been a major obstacle in this increasingly popular model system for coral-dinoflagellate endosymbiosis. Here we studied Aiptasia larvae feeding behavior and show its indispensability to trigger the first reports of settlement and metamorphosis in the system, finally closing the Aiptasia lifecycle. Surprisingly, the young gastrula-like planulae at just two days post fertilization actively feed on living crustacean nauplii, preferentially to heat-killed ones. Predation is dependent on functional stinging cells (nematocytes), indicative of complex neuronal control. Larvae fed daily dramatically increase in size and at 14 days post fertilization begin to morphologically change prior to settlement at high efficiency. Strikingly, dinoflagellate endosymbionts are neither required for larval growth nor measurably affect settlement dynamics, but are essential for spawning of the mature adult polyps. Our data show for the first time that gastrula-like planulae were capable of prey capture, suggesting carnivory in addition to filter feeding as a common strategy of this terminal life form. These data are discussed in the context of recent revelations on the evolution of basal metazoans.

## Introduction

In the nineteenth century, Ernst Haeckel proposed the developmental stage of the gastrula as the terminal life form of primitive multicellular first metazoans ^1^, an idea that became both influential and controversial ^2,3^. As the sister group to bilaterians, cnidarians are key to studying the evolution of development, and much work has been done to understand their wide variation of gastrulation modes using the key model anthozoan *Nematostella vectensis*, the starlet sea anemone (e.g. ^4^. Despite many advantages, *Nematostella* is not symbiotic with Symbiodiniaceae dinoflagellate algae, a trait found throughout many anthozoans and most famously in the ecologically critical reef-building corals. The small sea anemone *Exaiptasia pallida* sensu *diaphana* (commonly Aiptasia) has been established as a laboratory model for coral-algal symbiosis ^5,6^. Many resources have been developed in Aiptasia (Davy et al. 2012; Baumgarten et al. 2015), which can be reproducibly induced to spawn under laboratory conditions (Fig. 1A) ^7^. The developing embryo gastrulates by invagination (emboly) to form a planula larva, which after two days post-fertilization (dpf) has a spacious gastric cavity, a well-defined pharynx, nematocytes in the ectoderm, and an apical ciliary tuft (Fig. 1B-C) ^8^. Despite the relative ease of planulae larvae production in the laboratory, until now Aiptasia larvae have not been reported to undergo metamorphosis and settlement into primary polyps.

**Figure 1.**
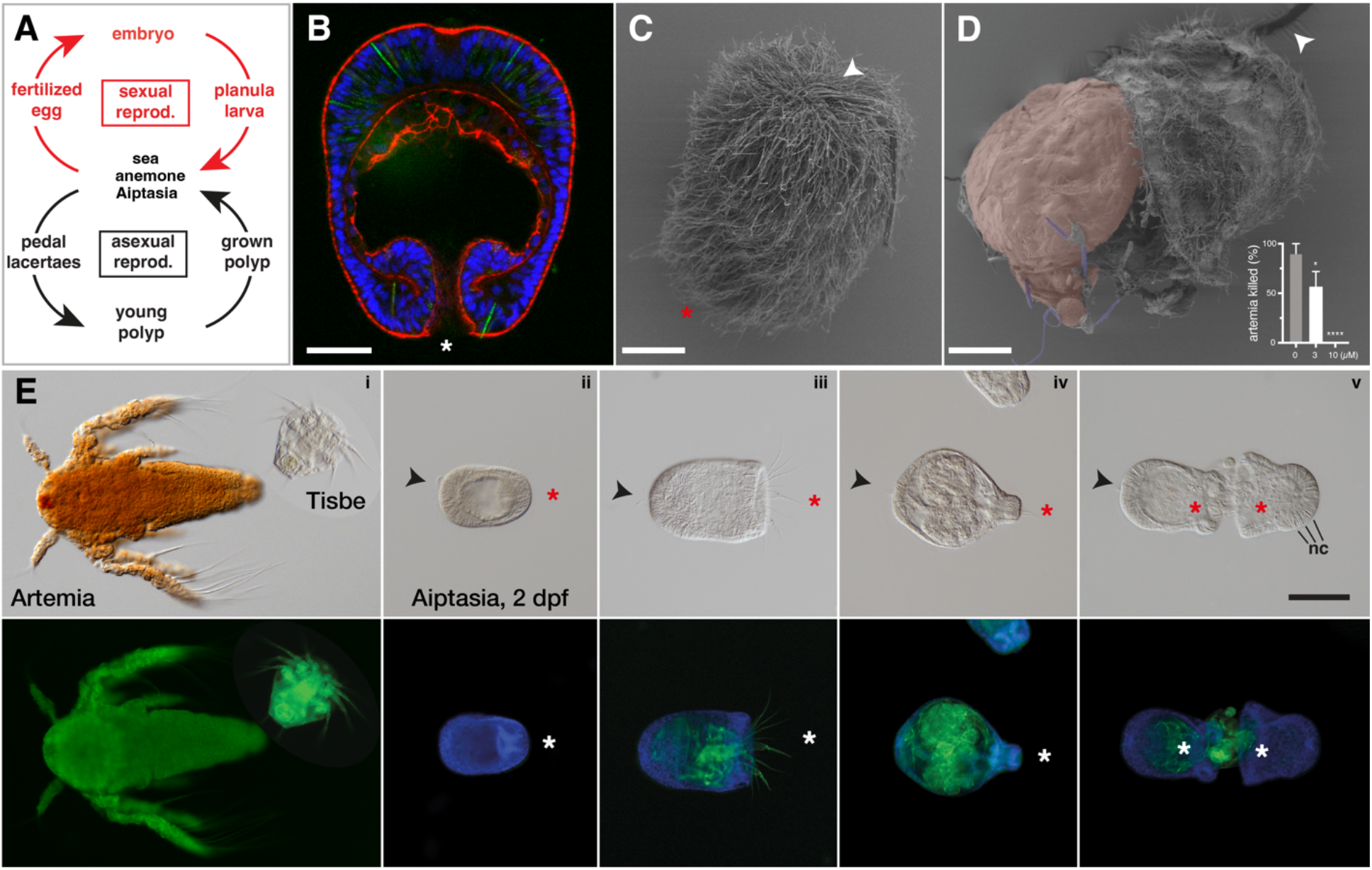
Carnivorous larvae of Aiptasia. (**A**) Life cycle of Aiptasia depicting asexual and sexual reproduction. (**B**) Representative images of a late gastrula / early planula larva at 2 days post fertilization (2 dpf; also in *E*). Blue, Hoechst; red, phalloidin; green, autofluorescence of nematocysts. The gastric cavity is lined by endoderm, particularly thickly toward the aboral end. The mouth is indistinguishable from the blastopore. Scale bar represents 25 μm. (**C-D**) Scanning electron micrographs of typical young Aiptasia planulae (3 dpf) showing extensive ectodermal surface cilia (*C*) and the ingestion of a *Tisbe* nauplius (*D*). The nauplius carapace (red) is penetrated by harpoon-like nematocysts (blue) from the larval oral end, and nematocyst activity is impaired with increasing concentrations of a small molecule inhibitor. The fraction of killed prey animals is presented as mean +/− S.D. from 3 independent experiments, analyzed by Student’s t-test. (*D*). Scale bar represents 25 μm. (**E**) Aiptasia larvae at 2 dpf and crustacean prey under DIC (*upper row*) and fluorescence (*lower row*) microscopy; blue, Hoechst; green, autofluorescence of chitin. Young Aiptasia planulae larvae require food of appropriate size like *Tisbe* nauplii, instead of the commonly used *Artemia* nauplii (*i*). *Tisbe* nauplii are predated by one (*iii, iv*) or multiple larvae (*v*) and completely ingested into the gastric cavity (*iii*-*iv*), which is then closed by the pharynx (*iv*). Scale bar represents 100 μm. (B-E) Asterisks, mouth; arrows, aboral apical tuft; nc, nematocysts.

While some triggers for cnidarian larvae settlement are chemical or other environmental cues ^9^, we focused on the dual nutrition sources of diet and algal symbionts, because Aiptasia eggs have low yolk content compared to *Nematostella* and *Acropora* (albeit an advantage in imaging studies). Similar to the planulae larvae of *Acropora* coral, *Fungia* coral, and *Anthopleura* anemones ^10,11^, Aiptasia planula larvae react to the presence of animal homogenate in the environment and can use mucus trails to ingest material, including inert particles, non-symbiotic algae, and symbiotic Symbiodiniaceae algae ^12,13^. The planula larvae establish robust symbiosis with various Symbiodiniaceae strains, which multiply in the larval endoderm over time ^8,13^. However, previous efforts of symbiont provision as well as exhaustive testing of various food sources did not lead to noticeable growth or settlement of larvae (our and others’ unpublished observations).

Here we show that Aiptasia larvae actively feed as early as two days after fertilization on live nauplii from the copepod *Tisbe* ^14,15^. This led to substantial growth of the larvae and, in a first report, to eventual settlement and metamorphosis. Endosymbionts are not necessary for growth, settlement, and metamorphosis in larvae, but rather exclusively for gametogenesis in adult Aiptasia animals and thus for successfully rearing polyps over several generations. Our data show for the first time that these cnidarian late gastrulae / early planulae were capable of prey capture, suggesting carnivory rather than filter feeding as a common strategy of this life form. This has implications for both the role of a predatory gastrula in the early evolution of (eu-)metazoans as well serves as breakthrough in the increasingly popular Aiptasia model system.

## Results

### Carnivorous feeding and growth of planula larvae

Surprisingly, we found that even early planulae at the gastrula stage (2 dpf; Fig. 1B) were able to hunt and catch freshly hatched *Tisbe* nauplii, which are smaller than the nauplii of the common food shrimp *Artemia salina* (Fig. 1C, D, E). *Tisbe* nauplii are caught by the Aiptasia larvae’s nematocysts (Fig. 1B, D), which penetrate the integument of the prey. *Tisbe* are then ingested into the planula gastric cavity through a widely opened mouth that subsequently closed for digestion (Fig. 1D, E; Video S1, S2), with undigested food expelled after several hours. Larvae fed with *Tisbe* nauplii grew continuously and dramatically in size, followed by eventual metamorphosis and settlement (Fig. 2A, B). In contrast, unfed larvae developed as described previously ^8^, i.e. total size and endoderm thickness steadily shrank over time (Fig. 2B). Live *Tisbe* nauplii were preferred to heat-killed nauplii by the Aiptasia larvae (Fig. 2B), and after 8 days of daily feeding, larvae grew large enough to also hunt and ingest Artemia. Inhibition of nematocyst discharge by a previously described small molecule inhibitor ^16^ prevented prey capture and led to increased prey survival (Fig. 1D).

**Figure 2.**
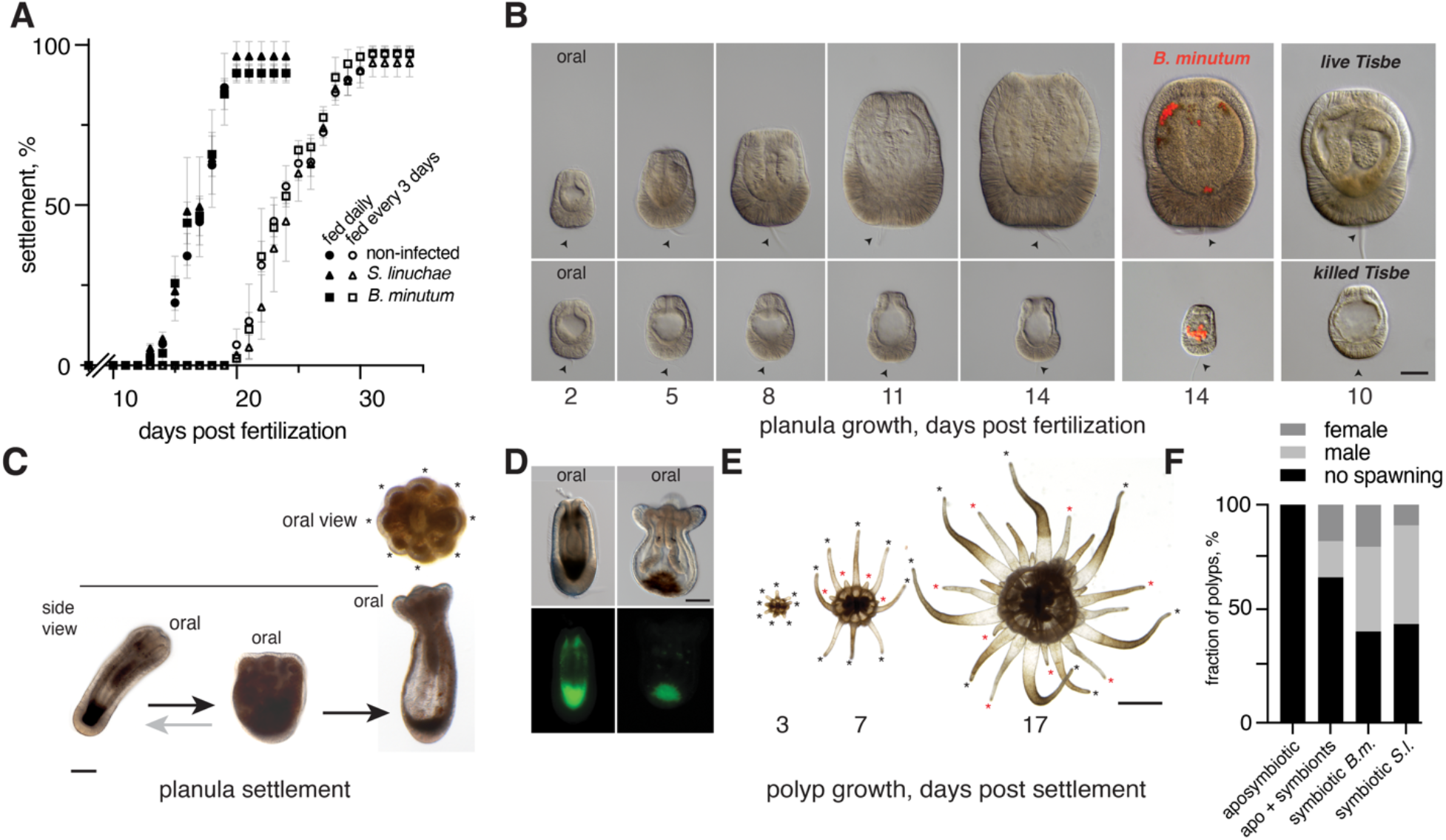
Larval and polyp development of Aiptasia. (**A**) Settlement dynamics of *Tisbe* nauplii-fed planulae larvae and larvae hosting either Symbiodiniaceae symbiont strain. Shown are average and standard deviation of 4 replicate experiments of 72 larvae per condition (experimentally lost larvae are not considered). No unfed larvae (0 of 144) underwent metamorphosis or settlement in any tested condition. (**B**) Growth of planulae and symbiotic planulae in fed-daily and unfed conditions over 14 days post fertilization (dpf), and comparison to larvae with Symbiodiniaceae symbionts and larvae fed with heat-killed *Tisbe* nauplii. Scale bar represents 50 μm. (**C**) Settlement of the planula larva and primary polyp, with cycling between elongated and laterally flattened forms before final settlement in latter form. Arrows indicate aboral apical tuft. Scale bar represents 100 μm. (**D**) Autofluorescence in live Aiptasia planulae appears after approx. 5-8 dpf and is brightest around settlement. Scale bar represents 100 μm. (**E**) Tentacle formation in growing polyps from 3-17 days post settlement (dps). Asterisks indicate the waves (sequence) of tentacle formation, with black as first (symmetric) wave primary polyp, red as second (asymmetric) wave. Scale bar represents 1 mm. (**F**). Sexual reproduction via spawning in settled Aiptasia primary polyps carrying symbionts (Sym), without symbionts (Apo), or aposymbiotic polyps infected with Symbiodiniaceae strains.

### Settlement of planula larvae

After approx. 14 days of daily feeding with *Tisbe* nauplii, planulae behavior began to markedly change as a precedent to settlement, the first report of which in Aiptasia. In addition to slower swimming, as in *Acropora* coral planulae (Grasso et al., 2011), Aiptasia planulae exhibited cycles of longitudinal lengthening and contraction together with substrate exploration with the aboral apical tuft (Fig. 2C; Video S3), concurrent with the novel appearance of autofluorescence at the aboral end (Fig. 2D). Settlement and metamorphosis occurred between 13 - 20 dpf (Fig. 2A), during which time planulae attached to the substrate at the aboral end, flattened, and displayed both mesenteries and 8 tentacle primordia (anlagen) (Fig. 2C; Video S3). Under a different dietary regime of feeding every third day, onset of settlement was delayed although the kinetics and final settlement efficiencies were strikingly similar, suggesting a size tipping point past which settlement was nearly always triggered (Fig. 2A).

### Effect of symbionts on planulae growth and settlement

We then sought to investigate whether Symbiodiniaceae algae symbionts affected settlement dynamics. Because the parental lines harbor either *Symbiodinium linuchae* strain SSA01 alone (male line CC7) or in combination with *Breviolum minutum* strain SSB01 (female line F003) ^7,17^, we infected larvae at 2 dpf with either SSA01 or SSB01 under the two *Tisbe* feeding regimes (Fig. 2A, B). All infected larvae displayed settlement dynamics indistinguishable from non-infected (aposymbiotic) larvae: only larvae fed with *Tisbe* nauplii grew and eventually settled, regardless of symbiotic status (Fig. 2A, B).

### From primary polyp to sexually mature polyp

Immediately after settlement, Aiptasia primary polyps grew and developed remarkably fast to reproductive maturity, and nutrition and symbiotic state affected both development and asexual and sexual reproduction differently. Most primary polyps displayed eight primary tentacle primordia that were radially symmetric, which shifted towards bilateral symmetry in the next tentacle wave around 7 days post settlement (dps), before eventually displaying sixfold symmetry after multiple tentacle waves (Fig. 2C, E). Growth patterns were identical in apo- and symbiotic primary polyps when fed at least twice weekly with *Artemia* nauplii, whereas less frequent feeding revealed an advantage in symbiotic polyps. Asexual reproduction by pedal laceration (Fig. 1A) was evident as early as 14 dps, and was similar in both apo- and symbiotic primary polyps.

Sexual reproduction in primary polyps could be induced ^7^ already at 6 months post-settlement, representing the first reports of F1 and F2 generations in Aiptasia (Fig. 2F). In contrast to asexual reproduction, sexual reproduction relied on symbiotic state: over half of symbiotic polyps hosting either algal strain spawned in one or two induction cycles (41 of 73), whereas no aposymbiotic polyps spawned (0 of 18). Critically, aposymbiotic polyps infected with symbionts recovered the ability to spawn (Fig. 2F). Finally, we combined gametes from spawned primary polyps to generate viable F2 planulae larvae, as well as back-crossing with parental lines in both sex combinations; all produced viable planulae larvae that could undergo settlement after feeding with *Tisbe* nauplii.

## Discussion

### Aiptasia model system

Our data provide clear evidence that feeding by the carnivorous predatory gastrula is the essential and limiting step in metamorphosis and settlement of Aiptasia planula larvae. This presents a major breakthrough, as Aiptasia continues to increase in popularity as a model system not only for coral-algae symbiosis but for other fields from embryology to ecotoxicology. It opens the door to functional genetics and manipulation in the amenable larvae phase ^18^ that can now be tested in settled adult polyps. Quick development of primary polyps to asexual competence allows expansion of manipulated clonal lines, which can also be sexually crossed or back-crossed through generations for the propagation of stable transgenic lines, thereby truly bringing Aiptasia and its toolkit into the molecular age.

### Role of symbiosis

Our results demonstrate that, surprisingly, Symbiodiniaceae algal symbionts do not have any observed effect on growth and settlement of Aiptasia planulae, through either signaling or nutritional contribution. The importance of diet over symbiosis appears to persist during primary polyp growth and development and into early asexual reproduction. However, in adult polyps the nutritional balance shifts towards the primacy of the symbiosis ^19,20^, including our observation of a striking reliance on symbiosis for Aiptasia sexual reproduction. This has been observed in the *Hydra*/*Chlorella* symbiosis ^21^, and although some aposymbiontic corals can spawn ^22^, the breakdown of symbiosis likewise rather severely affects sexual reproductive success in corals (e.g. ^23^)

### Evolutionary implications of the carnivorous gastrula

The carnivorous larvae of Aiptasia is remarkable for its ability to hunt live food only 2 dpf as a late gastrula / early planula, whereas most scleractinian corals develop into planulae after 3 or 4 dpf followed by settlement spontaneously or influenced by environmental chemical cues ^9-11^. The quick development and voracious appetite of Aiptasia is consistent with its lack of lipid-rich yolk in the embryo compared to fellow cnidarians. This lifestyle is likely an advantage in the heavily human-impacted environments in which Aiptasia are found worldwide, including aquaria and marinas ^24^, as they are comparatively eutrophic. Symbiosis establishment is nevertheless eventually critical in ontogeny, and the planulae autofluorescence we observed is consistent with the ‘beacon’ hypothesis of symbiont attraction in juvenile corals ^25^. However, the predatory life style might have been also an ancestral feature of the cnidarian gastrula. Derived cnidarians such as the hydrozoans do not develop a planula with functional mouth, and their yolk-rich embryos might have been an adaption to benthic life style. Taken together, the carnivorous gastrula of *Aiptasia* carries broad implications for the evolution of early emerging metazoans. Recent evidence supports the sister clade to all metazoans to be ctenophores ^26^, suggesting secondary loss in sponges. Our data are consistent with this pattern, as a putative shared origin between the ctenophores and planulazoans (i.e., cnidarians and bilaterians) of a nervous system and capturing cells ^27^, as well as, newly considered, a functional gut that allows active hunting in the voraciously predatory early gastrula larvae. Thus, the life style of the early gastrula using neurosecretory-related extrusive or adhesive cell types (i.e., nematocytes or cnidocytes in cnidarians and colloblasts in ctenophores) for predation ^28, 29^ might have driven the evolution of an organized nervous system in animals.

## Materials and Methods

### Organism culture

Aiptasia were cultured and induced to spawn as described ^7^ with the following modifications: animals were kept at 25°C in artificial seawater (ASW) made with Tropic Marin Pro Reef prepared in MilliQ water. Adult polyps were fed twice weekly with freshly hatched *Artemia salina* brine shrimp nauplii, followed by water exchange 2-5 h after feeding and weekly cleaning. Aiptasia larvae were collected and filtered as previously described ^7^. *Tisbe* copepods (planktino.de) were kept at a density of 20-30 animals per ml in ASW under aeration at 22-24°C. Cultures were dosed three times weekly with a nutrient solution (7.5% w/v in water) consisting of spirulina and spinach powders and baker’s yeast or Rotiwonder powder (planktino.de). *Tisbe* cultures were cleaned via ASW replacement while collecting adults in a 100 μm plankton sieve, and nauplii were further separated with a 60 μm plankton sieve. Clonal and axenic Symbiodiniaceae strains *Breviolum minutum* SSB01 (28S LSU identification at GenBank Accession MK692539) and *Symbiodinium linuchae* SSA01 (GenBank MK692538) were cultured as described ^13,30^.

### Aiptasia larvae feeding

Larvae were kept in 24-well plates with one larva per well in 1 ml ASW. Where applicable, larvae at 2 dpf were infected with the Symbiodiniaceae strains at a final concentration of 100,000 algal cells per ml. After 24 h of infection, larvae hosting at least one algal cell were kept in the experiment. Larvae were fed with 5-10 freshly hatched *Tisbe* nauplii daily or every three days starting at 2 dpf, and uneaten *Tisbe* were removed the following day. Larvae were transferred to fresh plates after approx. 2-3 feedings. Starting at 8 dpf for daily fed larvae, and 15 dpf for larvae fed every three days, larvae were additionally fed with 3-5 freshly hatched Artemia naupliI, and uneaten Artemia were removed the day after feeding. Settlement was documented daily. To test heat-killed food, larvae were fed with 5 live or 5 heat-killed (70°C for 10 min) *Tisbe* nauplii daily from 2 dpf to 9 dpf.

### Inhibition of nematocyst discharge

50 larvae (unfed, 4 dpf) were transferred to each well of a 12 well plate. The [2.2]paracyclophane compound 1 previously described to inhibit nematocyst discharge in *Hydra* ^16^ was added to the wells in triplicates at the indicated concentrations. The solvent DMSO was used in the control condition and the larvae were incubated for 30 min at 25°C. Then 10 artemia were added to each well and after 1 h the fraction of killed artemia per well was recorded.

### Staining, fixation, and imaging

For DIC and epifluorescence microscopy, larvae were stained with Hoechst 33258 for 1 h before fixation where applicable. Fixation was conducted 30-70 min or 24 h after feeding to document *Tisbe* uptake or larval growth, respectively, in 3.7% formaldehyde or 4% paraformaldehyde in ASW for 30 min at room temperature, followed by 3 washes in 0.05% Triton X-100 in PBS. After a final wash in PBS, larvae were embedded in glycerol on glass slides with glass coverslips. For live imaging of autofluorescence, animals were anesthetized in 40 mM chloral hydrate in ASW (larvae) or 180 mM MgCl2 (polyps) on glass slides with glass coverslips. Elongated larvae and top-view polyps were imaged live in ASW. For live videos, larvae and *Tisbe* nauplii were kept in 1 mm or 4 mm diameter wells made of Silicone Isolators™ Material (Grace Bio-Labs) in ASW on glass slides with a glass coverslip. Larvae, *Tisbe* and Artemia nauplii were imaged with a Nikon Eclipse 80i microscope using Differential Interference Contrast (DIC) or Fluorescence with a Nikon 10 x Plan Apo dry lens or a Nikon Plan Fluor 20× dry lens and a DS-Fi3 camera (Nikon Instruments). Oral views of polyps were captured using a Nikon SMZ25 with a SHR Plan Apo 0.5 x nosepiece and a DS-Ri2 camera. Videos were imaged using a Nikon Eclipse 80i microscope using Differential Interference Contrast (DIC) with a Nikon 10 x Plan Apo dry lens and a DS-Fi3 camera (Nikon Instruments) or with an iPhone 13 mini through LabCam® from iDu Optics® (New York, NY, USA).

For confocal microscopy (Fig. 1B), larvae were fixed at 2 dpf in 3.7% formaldehyde and 1x PHEM (60 mM PIPES, 25 mM 4-HEPES, 10 mM EGTA and 2 mM MgCl2, pH 6.9) in FASW for 30 min, followed by permeabilization in 20% DMSO and 0.1% Triton-X100 in PBS for 30 min. Larvae were then stained with Hoechst 33258 and Phalloidin Atto 565 for 4 h, washed thrice in 0.05% Tween20 in PBS, and mounted in 100% glycerol. Samples were imaged on a Leica SP8 laser scanning confocal at 63x magnification. For visualization of nematocyst autofluorescence, the detector for the 488 nm laser was set to reflection, and 3.5 μm stacks (7 z-planes) were recorded. Images were prepared using averaged Z-projections in ImageJ^31^.

For scanning electron microscopy, larvae were fixed in 2.5% glutaraldehyde in PBS for 2 h at 4 °C, rinsed in 0.1M cacodylate buffer pH 7.2, and post-fixed in 1% Osmium tetroxide in 0.1 M cacodylate buffer for 1 h at room temperature. Samples were rinsed in water and dehydrated in steps through a series of 50 to 100% acetone. Larvae were critical-point dried in a Leica CPD300 (Leica Microsystems, Vienna, Austria) and mounted on stubs with carbon adhesive discs. Finally, the larvae were sputter coated with 10nm gold/palladium and imaged with a Leo 1530 scanning electron microscope (Zeiss, Oberkochen, Germany).

### Induction of Aiptasia F1 spawning

F1 primary polyps were kept in groups of 15 in the same conditions as adult Aiptasia polyps above, and were fed twice weekly with Artemia nauplii for 5 months before exposure to spawn induction cues ^7^. Polyps were then kept individually and the identity of spawned gametes was documented.

## Acknowledgements

Supported by the DFG with grants to T.W.H. (SFB1324 A5 and D.A.C.H), to S.Ö. (SFB1324 B7 and Oe416/8-1) and through H2020 European Research Council (ERC Consolidator Grant 724715) to AG. Electron microscopy was performed by Charlotta Funaya and Larissa Eis at the Electron Microscopy Core Facility of Heidelberg University, and we are grateful for their technical assistance. We also thank Ulrike Engel and Nico Dross (Nikon Imaging Center of Heidelberg University) for their support in live cell imaging. The nematocyst inhibitor compound was kindly provided by Stefan Braese (KIT, Karlsruhe).We also thank Grisha Genikhovich (Vienna University) for his critical and helpful comments.

## Supplementary data

**Video S1** Hunting and feeding of Aiptasia I. Gastrula-like Aiptasia larvae, 2 dpf old, hunting on *Tisbe* nauplii (Interference contrast, real time).

**Video S2** Hunting and feeding of Aiptasia II. Feeding and uptake of a *Tisbe* nauplius by an Aiptasia larva, 2 dpf old (Interference contrast recording ca 60 minutes, time lapse 20X).

**Video S3** Settlement of Aiptasia planulae. Fed larvae of Aiptasia, 15 dpf, slow down swimming and elongate before they finally contract and adhere on the substrate to metamorphose into a primary polyp (Interference contrast, total recording time ca 6 minutes, time lapse 2X and real time).

